# Preparatory Suppression and Facilitation of Voluntary and Involuntary Responses to Loud Acoustic Stimuli in an Anticipatory Timing Task

**DOI:** 10.1101/2020.07.02.184143

**Authors:** An T. Nguyen, James R. Tresilian, Ottmar V. Lipp, Le-Anne Jacobs, Welber Marinovic

**Author notes:** Corresponding authors: An T. Nguyen, Welber Marinovic.

## Abstract

In this study, we sought to characterise the effects of intense sensory stimulation at different stages of preparation for an anticipated action on voluntary and involuntary behaviours. In our experiment, we presented unexpected loud acoustic stimuli (LAS) at four critical times during movement preparation to probe the state of the nervous system (Baseline, −1192 ms, −392 ms, and 0 ms relative to expected movement onset), and measured their effect on voluntary and involuntary motor actions (finger-press and eye-blink startle reflex, respectively).

Voluntary responses were largely facilitated by the LAS, leading to earlier and more forceful responses compared to Control and Baseline conditions. Notably, voluntary responses were significantly facilitated on trials where the LAS was presented early during preparation (−1192 ms). Eye-blink reflexes elicited by the LAS at −392 ms were significantly reduced and delayed compared to other time-points, indicating suppression of sub-cortical excitability. Despite being in a suppressive state, voluntary responses on these trials were still facilitated by the LAS.

The results provide insight into the mechanisms involved in preparing anticipatory actions. Induced activation can persist in the nervous system and can modulate subsequent actions for a longer time period than previously thought, highlighting that movement preparation is a continuously evolving process that is susceptible to external influence throughout the preparation period. Suppression of sub-cortical excitability shortly before movement onset is consistent with previous work showing corticospinal suppression which may be a necessary step before the execution of any voluntary response.

## 1 Introduction

The presentation of a loud acoustic stimulus (LAS) during movement preparation can trigger the release of a prepared response: the StartReact effect (Valls-Solé, Rothwell, Goulart, Cossu, & Munoz, 1999; Valls-Solé et al., 1995). This facilitatory effect occurs when participants are instructed to respond to the acoustic stimulus itself — in simple reaction time (RT) tasks where the imperative stimulus can be soft or loud (Honeycutt, Kharouta, & Perreault, 2013) — and also when participants are instructed to ignore these sounds — in anticipatory timing tasks where participants are instructed to respond at a fixed time after a warning cue (Carlsen & Mackinnon, 2010; Marinovic, Cheung, Riek, & Tresilian, 2014; Marinovic, de Rugy, Riek, & Tresilian, 2014; Tresilian & Plooy, 2006). In this study, we were interested in examining the time course of effects induced by the LAS during preparation for an anticipatory action. In particular, we sought to characterise the effects of the LAS on voluntary (finger-press) and involuntary responses (eye-blink startle reflex).

Kumru and Valls-Solé (2006) sought to determine whether the excitability of pathways that mediate the eye-blink reflex would be facilitated by the progressive increase in corticospinal excitability that occurs around 100 ms before movement onset (Chen, Yaseen, Cohen, & Hallett, 1998; Chye, Riek, de Rugy, Carson, & Carroll, 2018; Ibanez, Hannah, Rocchi, & Rothwell, 2019; Leocani, Cohen, Wassermann, Ikoma, & Hallett, 2000; MacKinnon & Rothwell, 2000) by delivering a LAS together with or after (20 to 100 ms) the visual go-stimulus. They showed that the timing of the LAS in relation to the go-stimulus had no clear effect on the amplitude of the eye-blink reflex, suggesting that movement preparation either has no effect over sub-cortical circuits or is at a constant level over the short time window tested. Similarly, Marinovic, de Rugy, Lipp, and Tresilian (2013) examined the amplitude and timing of the eye-blink reflex while delivering the LAS before and after the go-stimulus (−65 to 105 ms). The results showed a suppression in blink amplitude when the LAS was presented after the go-signal (Experiment 2) but not before (Experiment 1), suggesting that the associated sub-cortical circuits were inhibited shortly after (65 ms) the presentation of the go-stimulus, as opposed to being facilitated. These latter results are in contrast to those by Lipp, Blumenthal, and Adam (2001) who observed a facilitation in blink amplitude at a similar interval (60 ms post-stimulus) when participants were asked to perform a discrimination task (see also Lipp, Alhadad, & Purkis, 2007). One difficulty with interpreting these differing results is that these tasks typically involve the presentation of discrete visual and acoustic stimuli (e.g. visual go-stimulus and LAS), which depending on the stimulus onset asynchrony (SOA) could lead to modulation of the eye-blink reflex even in the absence of movement preparation (Boelhouwer, Teurlings, & Brunia, 1991; Burke & Hackley, 1997). To circumvent this problem, we employed an anticipatory timing task where the only stimulus presented discretely during a trial was the LAS. The advantages of this task are twofold: first, it can be reliably used to test the triggering of motor actions by LAS (Marinovic, de Rugy, et al., 2014; Tresilian & Plooy, 2006), and second, we and others have described the time course of corticospinal excitability of task relevant and irrelevant muscles using this task (Ibanez et al., 2019; Marinovic, Flannery, & Riek, 2015; Marinovic, Reid, Plooy, Riek, & Tresilian, 2011).

Most studies in the StartReact literature have used RT tasks with variable foreperiods, and it is clear from the available data that the predictability of the go-stimulus does impact the StartReact effect (see Leow et al., 2018). Some studies, however, have used tasks with greater temporal predictability. For example, using a fixed foreperiod (3.5 seconds) RT task, MacKinnon, Allen, Shiratori, and Rogers (2013) showed that the contingent-negative variation (CNV) measured from the electroencephalogram (EEG) become more negative seconds before the time of movement onset, and the more negative the CNV was, the larger the probability of observing an early release of the prepared action by a LAS. Marinovic et al. (2013) showed a linear decrease in RT and an increase in response vigour as the LAS was presented later during an RT task with a predictable foreperiod (1 second). Carlsen and Mackinnon (2010) found that most responses cannot be initiated early — defined as a response that occurs within 250 ms after an acoustic stimulus — by a LAS presented between 1500 to 500 ms before the expected time of movement onset in anticipatory timing tasks (see also Carlsen, Chua, Timothy Inglis, Sanderson, & Franks, 2008). However, the probability of releasing a response earlier than normal increases dramatically when the LAS is presented later in the trial (within 150 ms prior to movement onset time). Our group has employed anticipatory timing tasks to examine movement preparation in past research and consistently reported both an early release of the prepared actions and an increase in response vigour (Marinovic et al., 2015; Marinovic, Tresilian, de Rugy, Sidhu, & Riek, 2014; Tresilian & Plooy, 2006), but the time course of these effects has not been explored over longer time windows (> 200 ms before the expected time of movement onset). In particular, it is unknown whether LAS presented more than a second before the expected time of movement onset in an anticipatory timing action can affect the execution of voluntarily prepared actions, modulating their RT and vigour.

In the present study, we sought to investigate how the delivery of a task-irrelevant LAS at different times during movement preparation for anticipatory actions can affect the manifestation of voluntary and involuntary responses. We presented LAS at four critical times during the course of a trial: 1) Before trial onset (Baseline measurement), 2) immediately after a cue started moving towards an intersection zone (at trial onset), 3) before corticospinal excitability was expected to rise above baseline levels (during preparatory corticospinal suppression, see (Ibanez et al., 2019; Marinovic et al., 2011), and 4) when the cue intercepts with the stationary target (expected time of movement onset). We hypothesized that the sub-cortical excitability would reflect the pre-movement suppression that occurs between 500 and 200 ms before movement onset (Ibanez et al., 2019; Marinovic et al., 2011), that is, longer latencies and reduced amplitude of the eye-blink reflex when the LAS was presented about 400 ms prior to the expected time of movement onset but not after or during baseline measurements. We also expected that voluntary responses would be released earlier and more forcefully even if LAS were delivered during corticospinal suppression. Lastly, we predicted that the LAS at trial onset would not affect the timing of voluntary responses but would facilitate the eye-blink reflex given the short latency between visual motion onset and LAS (Neumann, Lipp, & Pretorius, 2004).

## 2 Method

### 2.1 Participants

Twenty-three participants consisting of undergraduate psychology students and volunteers from Curtin University were recruited for this study. One participant was excluded for low performance on Control trials, and five were excluded for a low number of blink trials (fewer than 2 accepted blinks per condition). The final sample consisted of 17 [*M*(*SD*)_*ase*_ = 21.65(3.72) years, Range = 17-33 years, 13 Females], All participants were right-handed, reported having normal or corrected vision, and to having no diagnosed or known neurological conditions. All participants provided written informed consent before starting the experiment and the protocol was approved by the human research ethics committee of Curtin University (Approval Code: HRE2018-0257).

### 2.2 Anticipatory Timing Task

As a brief overview of the task, participants were instructed to synchronise the onset of their response witłi tłie intercept of two rectangles by pressing on a force sensor with their right index-finger (See Figure 1). On 20% of trials, a LAS was pseudo-randomly presented at different times during before the intercept. Participants were simply instructed to ignore this sound and respond to the task as per usual.

**Figure 1:**
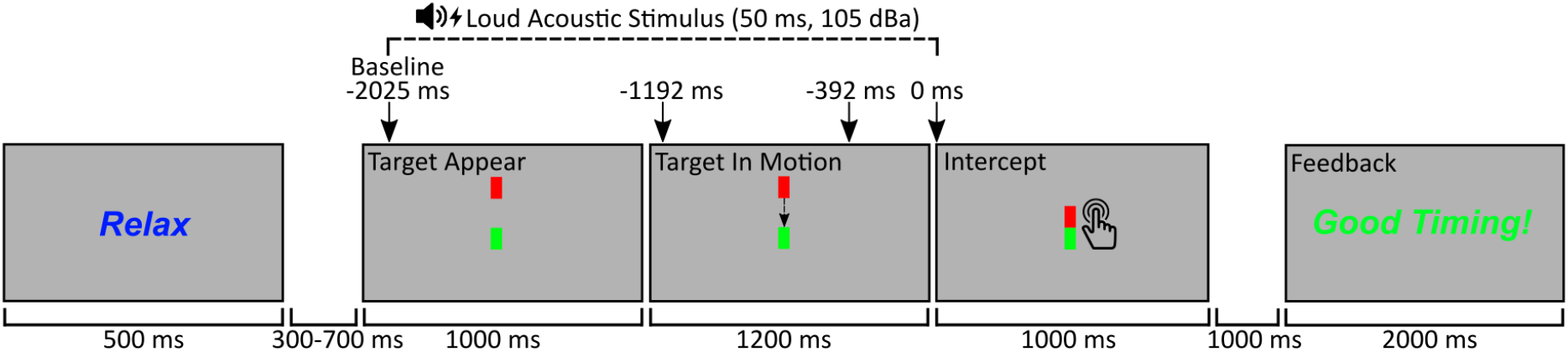
Diagram depicting the sequence of trial events. Participants were instructed to synchronise their response with the intercept of the two rectangles. On 20% of trials, a Loud Acoustic Stimulus (LAS) was randomly presented at one of four positions during the foreperiod (Baseline/-2025 ms, −1192 ms, 392 ms or 0 ms relative to intercept). Participants were instructed to ignore the LAS and respond to the intercept

The task was presented using MATLAB 2015b and Psychtoolbox version 3.0.11 (Brainard, 1997; Kleiner et al., 2007; Pelli, 1997) on a 22 inch Samsung LCD monitor (Model: 2233RZ, 1920 x 1080 resolution, 120 Hz refresh rate). Participants were seated 70 cm away from the monitor. Before the experiment, participants were presented with on-screen and verbal instructions, with demonstrations of the trial sequence and the LAS. They then completed a practice block consisting of 12 trials in a fixed sequence with one LAS trial, followed by a single experimental block consisting of 200 trials (160 Control trials, 40 LAS trials).

On each trial, ‘Relax’ was presented for 500 ms, followed by a blank screen of variable duration (300-700 ms). A red and green rectangle (4×9 mm) were then presented at the top and centre of the display, static for 1000 ms. Following this, the red rectangle descended towards the green rectangle, in motion for 1200 ms and stopping when the two edges touch. On LAS trials, a LAS was pseudo-randomly presented at one of four different time-points; 1) during the static period 175 ms after the rectangles appeared or −2025 ms from intercept (Baseline), 2) 8 ms after the target began moving or −1192 ms from intercept, 3) when the target was close (−392 ms) to the intercept, and 4) together with the intercept. The pseudo-randomisation occurred such that a LAS would not be presented on two successive trials. The LAS was a 50 ms burst of broadband white-noise at an intensity of 105 dBa, generated using an external custom-made white-noise generator, directly connected to high fidelity stereophonic headphones for low-latency presentation (Seinheiser model HD25-1 II, frequency response 16Hz to 22 kHz; Sennheiser Electronics GmbH & Co. KG, Wedemark, Germany). Sound intensity was measured with Briiel and Kjacr sound level meter (type 2205, A weighted; Briiel & Kjacr Sound & Vibration Measurement, Naerum, Denmark) placed 2 cm from the headphone speaker. Stimulus rise-time was measured to be 1.25 ms from the headphones.

Participants were instructed to time the onset of their response with the intercept of the two rectangles. Participants responded by quickly and briefly pressing their right index-finger on a force sensor (SingleTact, Model: CS8-10N) embedded in a shell resembling a computer mouse. Feedback was provided at the end of each trial 2000 ms after the intercept, for 2000 ms. If participants initiated their response within ± 30 ms of the intercept “Good Timing” was presented. Otherwise, “Too Early” or “Too Late” was presented if they responded outside the 60 ms response window. If no detectable response was on made on the trial, “No Response Detected” was presented. No feedback was presented on LAS trials and “Probe Trial” was presented instead.

### 2.3 Force and EMG Data Acquisition, Data Reduction and Measurement

Force data acquired from the force sensor were continuously recorded for the duration of the trial, digitised at 2000 Hz using a National Instruments data acquisition device (USB-6229 BNC multifunctional DAQ). The data were filtered using a low-pass second-order Butterworth filter with a cut-off frequency of 20 Hz. As our measure of response timing, we calculated movement onset time relative to the intercept from the tangential speed time-series derived from the force data using the algorithm recommended by Teasdale and colleagues (1993). Trials with onset latencies exceeding 150 ms before and after the intercept were excluded from further analysis. On average, 24.17 (*SD* = 38.53) trials per participant (12.08% of total trials) were rejected for invalid response times. We also measured peak force as our measure of response vigour.

We recorded EMG activity from the right Orbicularis Oculi muscle using a pre-amplified bi-polar set-up. We used 8 mm Ag/AgCl sintered electrodes, one electrode was placed under the pupil, the second was placed laterally and slightly higher than the first electrode, approximately 1 cm edge-to-edge and a ground electrode was placed on the right mastoid region. We used a Neurolog Systems Digitimer Pre-Amplifier (Model: NL82O) and Amplifier (Model: NL905), with a 50-1000 Hz pass-band filter and Gain set to 1000. These data were also digitised using the National Instruments DAQ.

The EMG data were processed offline using a semi-automatic procedure in R statistics and R studio. The data were down-sampled to 1000 Hz, rectified using the ‘rectification’ function in ‘biosignalEMG’ package (Guerrero & Macias-Diaz, 2018), and smoothed using 5-point moving average with the ‘rollapply’ function in the ‘zoo’ package (Zeileis, Grothendieck, Ryan, Ulrich, & Andrews, 2020). The Bonato, D’Alessio, and Knaflitz (1998) method was used to automatically detect the blink onset latency, using the ‘onoff_bonato’ function in the ‘biosignalEMG package’ (sigma n = 2 times the standard deviation of activity within 0-200 ms prior to the LAS). Multiple passes of this step were run. If no onset was detected, another pass was run with the threshold gradually increasing (by increments of 0.2 times baseline variability, 10 times) then decreasing (from 1 in increments of 0.2 times baseline variability, 2 times), until an onset was detected within 20-80 ms. We also measured baseline-to-peak EMG amplitude occurring after blink onset latency. Each trial was visually inspected, and corrections were made to onset and peak latencies where necessary. Acceptable onset latencies were within 20-80 ms from LAS onset. Responses outside this window were excluded from further analyses of blink data (Blumenthal et al., 2005). Trials with a flat EMG response were classified as ‘non-response trials’. Trials containing excessive noise, artifacts, or voluntary activation within 20 ms after LAS onset were classified as ‘missing’ trials. Non-response and missing trials were not included in further analyses of blink data. On average, 1.06 (*SD* = 1.98) trials per participant (2.65% of LAS trials) were rejected for invalid blink latencies, 3.47 (*SD* = 5.51) trials for non-response trials (8.68% of LAS trials), and 1.82 (*SD* = 1.46) trials for missing trials (4.56% of LAS trials). On average, 141.12 (*SD* = 31.82) Control trials (88.2%) and 28.59 (*SD* = 8.75) LAS trials (71.5%) were retained.

### 2.4 Statistical Analysis

Statistical analyses were conducted in R statistics and R studio. We used linear mixed model analyses using the “liner” function from the “LinerTest” package (Kuznetsova, Brock-hoff, Christensen, & Jensen, 2019) to examine the effects of the LAS on movement onset time, peak force, blink onset latency and blink amplitude using trial-level data. For movement onset and peak force, we modelled each variable with ‘Condition’ (Control, LAS at baseline/-2025 ms, −1192 ms, −392 ms, and 0 ms) as the fixed-effect predictor. For blink latency and blink amplitude, we modelled ‘LAS position’ as the fixed-effect predictor (LAS at baseline/-2025 ms, −1192 ms, −392 ms and 0 ms). For all models, participant intercepts were modelled as the random effect. We presented the results as F-values using the Satterthwaite’s approximation method (Satterthwaite, 1941), using the ‘anova’ function. We used the ‘r2beta’ function from the ‘r2glmm’ package to calculate effect size as Partial R^2^ (*R^2^_P_*) (Jaeger, 2017). For follow-up pairwise contrasts, we computed estimated marginal means using the ‘emmeans’ function from the ‘emmeans’ package (Lenth, Singmann, Love, Buerknerm, & Herve, 2020). We presented the results of these pairwise categorical comparisons as t-ratios (mean difference estimate divided by standard error) with degrees-of-freedom estimated using the Kenward-Roger method, and the Hochberg (1988) method was used to control for multiple comparisons.

## 3 Results

We found that the timing and force of the voluntary response differed significantly between Control trials and LAS presentations across the foreperiod (Movement Onset: F(4,2967) = 8.99, *p* < .001, *R^2^_P_* = .012 [.006 - .022]; Peak Force: F(4,2967) = 8.44, *p* < .001, *R*^2^_*P*_ = .011 [.006 - .021]). Follow-up pairwise contrasts for movement onset times showed that presenting the LAS while the target was in motion (−1192 ms and −392 ms) resulted in earlier onset times compared to Baseline and Control trials (except for −1192 ms *vs*. Baseline). When the LAS was presented at the intercept, onset times were not facilitated but instead showed signs of inhibition. The pattern of peak force results mirrored that of onset times, showing marked increases while the target was in motion, followed by a decrease when the LAS was presented at the intercept. Follow-up comparisons showed significantly larger peak force when the LAS was presented at −392 ms, compared to Control and Baseline trials.

The timing and magnitude of the eye-blink reflex elicited by the LAS also changed significantly during the preparation foreperiod (Blink Latency: F(3,570.11) = 5.27, *p* = .001, *R*^2^_*P*_ = .027 [.010 - .063]; Blink Amplitude: F(3,586) = 4.08, *p* = .007, *R*^2^_*P*_ = .021 [.006 - .054]). Follow-up pairwise contrasts showed significantly longer blink latencies at −392 ms prior to the intercept, compared to all other conditions. Blink amplitude results were consistent with blink latency findings showing significant reductions for LAS at −392 ms compared to the intercept (0 ms).

## 4 Discussion

Movement preparation is a dynamic process involving cortical and sub-cortical areas of the brain (Kaufman, Churchland, Ryu, & Shenoy, 2014; Paradiso, Cunic, & Chen, 2004; Paradiso, Cunic, Saint-Cyr, et al., 2004). It is well established that the presentation of a LAS can interact with these preparatory processes, decreasing movement onset times and increasing response vigour (Marinovic & Tresilian, 2016; Valls-Solé, 2012). Many studies have examined the effects of the LAS close to the time of response initiation (100 ms before or after response onset time) (Kumru & Valls-Solé, 2006; Marinovic et ah, 2013). However, the extended time-course of the interactions between LAS and movement preparation is largely unknown. Therefore, in the present study, we sought to fill this gap in knowledge about extended time course of preparation for anticipatory actions. We used LAS to probe the excitability of the nervous system at four critical time intervals from the expected response onset time (Baseline/-2025 ms, −1192 ms, −392 ms and 0 ms) and examined their effect on voluntary and involuntary responses.

### 4.1 Voluntary responses effects

In line with our predictions, we observed facilitation of the voluntary responses when LAS were presented at longer time periods (up to 1.2 seconds) from the intercept, while the target was in motion. However, voluntary responses were not facilitated when the LAS was presented 2 seconds prior, while the target was presented but was not in motion (Baseline condition). These findings indicate that increased activity induced by the LAS can persist for a prolonged period (1.2 seconds), but only when participants were to engage in movement preparation shortly. Response facilitation when the LAS was presented in early preparation was evident for both movement onset time and peak force. These findings are largely consistent with previous research showing early response initiation and increased forcefulness when the LAS is presented close to the time of movement onset (Marinovic et al., 2013). Similar to our study, Carlsen and Mackinnon (2010) used LAS in anticipatory tasks (clock condition), delivering the LAS at −1500 ms, −500 ms, and −150 ms before the expected time of response onset. They analysed the proportion of trials a response was produced within very short latency (< 250 ms) and found that LAS evoked fast responses were elicited in 0% of the trials at −1500 ms, 17.5% at −500 ms, and 100% at −150 ms. Their results were interpreted as evidence that movement preparation for anticipatory actions is delayed until < 500 ms before the expected time of movement onset, when continuous temporal information was provided. Our results, however, clearly indicate that the facilitatory effects of irrelevant sensory stimulation are retained in the circuits responsible for response preparation for more a much longer time (> 1 second).

The absence of movement facilitation on Baseline LAS trials (target stationary) could be due to it falling outside the target-movement period or its long temporal distance (−2025 ms) from the intercept. The former would suggest that the impact of LAS on the excitability of the system is state-dependent. LAS activation may persist for longer only if induced during the target-movement interval, where the system is in an active preparatory state, and decays faster if induced outside this interval. We have previously found state-dependent effects of LAS on the primary motor cortex during an anticipatory timing task (Marinovic, Tresilian, et al., 2014). The observed facilitation of the response when the LAS was presented at −1192 ms (175 ms after the target-movement), indicates that the active preparatory state can be rapidly initiated by visual cues.

Another interesting finding in our study was the inhibitory effect that the LAS had on response onset times at 0 ms - where response onset times were significantly delayed compared to control trials. Given the anticipatory nature of the task, it was expected that the LAS would have a negligible effect on response onset time because motor commands should have already been issued (see Tresilian & Plooy, 2006). This finding is intriguing but resembles those reported by Xu-Wilson, Tian, Shadmehr, and Zee (2011). In more detail, in their experiment 3, TMS was delivered just before participants initiated oblique saccades. They found that stimulation very close to saccade onset slowed or even paused their onset (< −40 ms to saccade onset), but stimulation at even earlier times could reduce saccade latency (see also Castellote, Kumru, Queralt, & Valls-Solé, 2007) — this effect was independent of the site of stimulation. Although response onset time to the LAS at 0 ms was still negative (See Figure 2), that is, participants responded on average before the expected time of movement onset. All that is needed to obtain such a result is that some responses, not all of them, be delayed for an increase in response latency to occur; visual inspection of the distribution of response onset times on Control trials indicated that 20.84% and 10.54% of responses occurred 20 and 40 ms after the intercept respectively, late enough to be affected by the LAS at 0 ms. Thus, our results indicate that the mechanism by which inhibition may have occurred at 0 ms in our study is the same as that observed by Xu-Wilson et al. (2011) and involves the startle circuit — reticular formation (Yeomans, Li, Scott, & Frankland, 2002). While modulation of timing was an unexpected finding, the LAS was still expected to increase peak force as the motor command issued by the motor cortex should still be unfolding (Stahl & Rammsayer, 2005). We did observe an increase in peak force above baseline levels at 0 ms, but this effect was not statistically significant and smaller than that observed at −392 ms.

**Figure 2:**
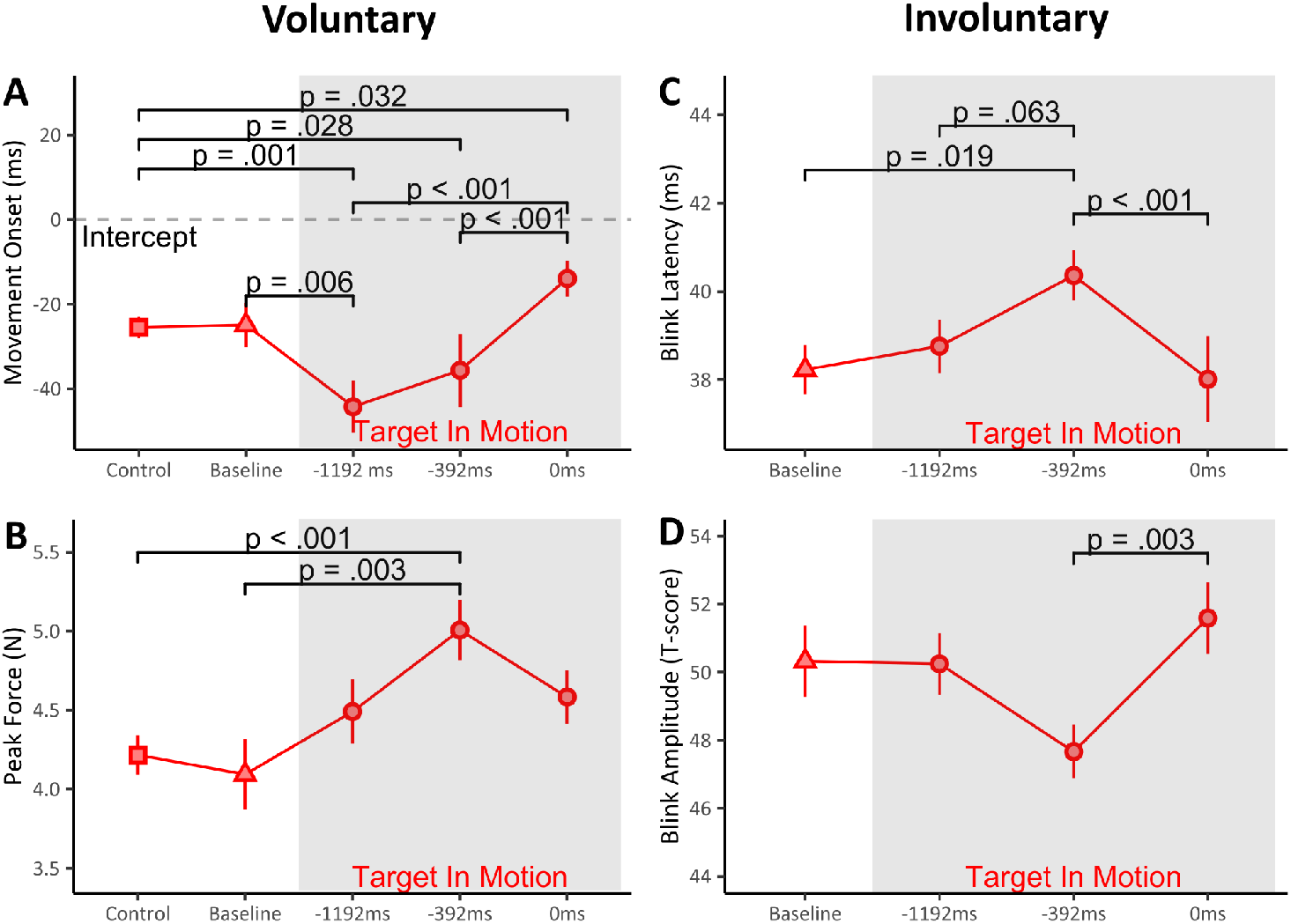
Grand-average values for (**A**) Movement Onset (relative to intercept), (**B**) Blink Latency, (**C**) Peak Force, and (**D**) Blink Amplitude across Control and LAS conditions (red, Baseline/-2025 ms, −1192 ms, −392 ms, 0 ms) with within-subjects standard error bars. The grey shaded area highlights the period when the red rectangle was descending towards the green rectangle. Significance brackets with p-values mark all marginally (i.e. p < .100) and statistically significant contrasts.

### 4.2 Involuntary responses effects

In comparison to voluntary responses, the blink reflex revealed a different pattern of modulation over the time course of preparation. Inhibitory effects on the eye-blink reflex emerged 392 ms before the intercept, in both latency and amplitude, reflecting suppression of sub-cortical excitability but not at any other timepoint. Despite the mixed findings in the literature (see Introduction), the clear suppression of the blink reflex we observed was not unexpected during late movement preparation. Anthony and Putnam (1985) had shown suppression of the blink reflex in a reaction time task and so did our group (Marinovic et al., 2013). What is intriguing, however, is that during early preparation (−1192 ms) we found no evidence of blink modulation, even though the LAS effect on the subsequent voluntary response was clear. That is, even though the state of preparation seemed to have started early in a trial, modulation of the blink reflex only occurred much later. The time of blink reflex suppression in our study aligns with results Ibanez et al. (2019) reported for pre-movement suppression of corticospinal excitability in an anticipatory timing task — between 500 to 200 ms prior to EMG onset — using transcranial magnetic stimulation (TMS): a phenomenon necessary for self-paced, anticipatory and reactive motor responses (see also Marinovic et al., 2011). Noteworthy, the sharp increase in blink amplitude and the reduction of response latency between the −395 ms and 0 ms timepoints is consistent with the removal of sub-cortical suppression during reaction time tasks (see also Lipp et al., 2007. Using transcranial electric stimulation (TES) of the motor cortex, Starr, Caramia, Zarola, and Rossini (1988) found that motor-evoked potentials were completely suppressed −80 ms or earlier before EMG onset, but became progressively larger from −60 ms until movement onset. TES activates the axons of corticospinal neurons directly, bypassing the soma (Di Lazzaro et al., 2004; Rothwell et al., 1994), indicating that the effect reported by Starr et al. (1988) reflects a release of sub-cortical inhibition connected to the normal transition from preparation to execution. Altogether, the direction and time of the effects of LAS on involuntary responses do not seem to be coincidental. That is, they appear to be mainly driven by voluntary movement preparation rather than an effect of attention being diverted to a different sensory channel (Brunia, 1993).

### 4.3 Conclusion

In the present study, we used LAS to characterise the extended time-course of changes in nervous system excitability during the preparation of anticipatory actions. Our data showed that LAS can have significant facilitatory effects on voluntary motor responses from up to 1.2 seconds after stimulation, much longer than previously shown. This result suggests that activation induced by LAS can remain in response-related circuits for extended periods, even when induced during very early phases of preparation. Additionally, we also found evidence of a transition from sub-cortical suppression to facilitation during mid-to-late stages of preparation in the involuntary response, as well as inhibitory-effects on the voluntary response to the LAS during the movement execution phase. Altogether, these findings support the view that accessory sensory stimulation can systematically interact with cortical and sub-cortical excitability during movement preparation. Understanding the nature — inhibitory *vs*. excitatory — and the extended time course of this interaction is important because recent findings indicate that there is a causal link between movement preparation and sensorimotor learning (Vyas, O’Shea, Ryu, & Shenoy, 2020), suggesting the type of accessory sensory stimuli we employed can serve as a tool for movement acquisition or rehabilitation.

## Funding Bodies

This study was supported by a Discovery Project grand from the Australian Research Council (DP180100394) awarded to W.M and O.V.L.

